# *In vivo* photopharmacology enabled by multifunctional fibers

**DOI:** 10.1101/2020.03.28.012567

**Authors:** James A. Frank, Marc-Joseph Antonini, Po-Han Chiang, Andres Canales, David B. Konrad, Indie Garwood, Gabriela Rajic, Florian Koehler, Yoel Fink, Polina Anikeeva

## Abstract

To reversibly manipulate neural circuits with increased spatial and temporal control, photoswitchable ligands can add an optical switch to a target receptor or signaling cascade. This approach, termed photopharmacology, has been enabling to molecular neuroscience, however, its application to behavioral experiments has been impeded by a lack of integrated hardware capable of delivering both light and compounds to deep brain regions in moving subjects. Here, we devise a hybrid photochemical genetic approach to target neurons using a photoswitchable agonist of capsaicin receptor (TRPV1), *red*-AzCA-4. Using the thermal drawing process we created multifunctional fibers that can deliver viruses, photoswitchable ligands, and light to deep brain regions in awake, freely moving mice. We implanted our fibers into the ventral tegmental area (VTA), a midbrain hub of the mesolimbic pathway, and used them to deliver a transgene coding for TRPV1. This sensitized excitatory VTA neurons to *red*-AzCA-4, and allowed us to optically control conditioned place preference using a mammalian ion-channel, thus extending applications of photopharmacology to behavioral experiments. Applied to endogenous receptors, our approach may accelerate studies of molecular mechanisms underlying animal behavior.

## INTRODUCTION

To illuminate the molecular mechanisms underlying neuronal function, a number of chemical and genetic techniques have been developed to place cells under optical control.^1–3^ As demonstrated by optogenetics, the enhanced spatiotemporal precision attained using an optical stimulus is enabling to understanding how specific circuits control behavior and animal physiology.^1,4^ However, optogenetics has so far relied on exogenously expressed microbial rhodopsins, thus offering a means to investigate electrophysiological, but not molecular, mechanisms underlying neural dynamics and behavior. Light-controllable ligands present an attractive alternative to optogenetics, as they add an optical switch to a pharmacologically-defined target.^5^ Of particular utility are azobenzene photoswitches, which can be synthetically incorporated into a small-molecule ligand.^6,7^ Their activity can then be reversibly tuned using different wavelengths of light *via trans/cis* isomerization. Termed photopharmacology,^8^ this approach has been applied to manipulate ion channels, G protein-coupled receptors (GPCRs), enzymes, components of the cytoskeleton, and even cellular membranes.^6,9–14^ These photoswitchable small molecules can target endogenous receptors, or alternatively, genetically inserted exogenous proteins to create a wide variety of signaling events. Unfortunately, the utility of photopharmacology for studying mammalian behavior has been hindered by the lack of available hardware for delivering the chemical probes and optical stimuli to non-superficial brain regions. Although several recent studies have applied azobenzene-based probes to control behavior in freely moving rodents, they required the use of separate cannulas and waveguides for ligand and light delivery.^15–17^ This adds surgical complexity and possibly contributes to tissue damage due to the device footprint. As such, the development of integrated hardware that allows both chemical and optical stimulation in deep brain regions in freely moving subjects may facilitate the adoption of photopharmacology by the neuroscience community.

In this study, we devise a photochemical genetic method to manipulate neural circuits using the photoswitchable capsaicin (CAP) analog, *red*-AzCA-4.^18^ To translate this approach from cell culture to behavioral experiments, we used thermal drawing to create a flexible multifunctional fiber that can deliver optical and chemical stimulation to deep brain regions in mice. We targeted the ventral tegmental area (VTA), a midbrain region involved in dopaminergic reward signaling, by injecting a virus coding for the transient receptor potential vanilloid family member 1 (TRPV1, capsaicin receptor). This sensitized excitatory VTA neurons to *red*-AzCA-4, which could manipulate preference behavior in freely moving mice in a light-dependent manner. This work sets the stage for applications of photopharmacology in manipulating neural circuits *in vivo* to unveil molecular mechanisms underlying behavior.

## RESULTS

### Optical control of TRPV1-expressing neurons using *red*-AzCA-4

We previously developed photoswitchable CAP analogs that can optically control TRPV1, a non-selective cation channel which is expressed primarily in nociceptive neurons.^19,20^ The first generation probe, AzCA-4, includes an azobenzene photoswitch that can be isomerized between the *cis*/*trans* forms with UV-A/blue light, respectively.^21^ AzCA-4 was more potent in the *cis*-form, and isomerization could reversibly control endogenously expressed TRPV1 in sensory neurons *in vitro* and *ex vivo*. However, due to the phototoxicity and poor tissue penetration of UV-A (350-380 nm) irradiation required to generate the *cis*-isomer, AzCA-4 was not suitable for applications *in vivo*.^21^ A red-shifted derivative, *red*-AzCA-4,^18^ which possesses four chlorine atoms at the *ortho*-positions of the azobenzene moiety (**Fig. 1a**), was designed to isomerize to the more potent *cis*-form with green (500-590 nm) as well as the UV-A light, while returning to *trans*-form upon exposure to blue light (410-480 nm). This increases its applicability towards experiments in intact tissue, as longer wavelength visible light is less phototoxic.^22,23^ Although *red*-AzCA-4 was shown to behave similar to AzCA-4 in cultured cells heterologously expressing TRPV1, its utility in modulating activity of TRPV1-expressing neurons *in vitro* and *in vivo* remained unexplored.

**Figure 1.**
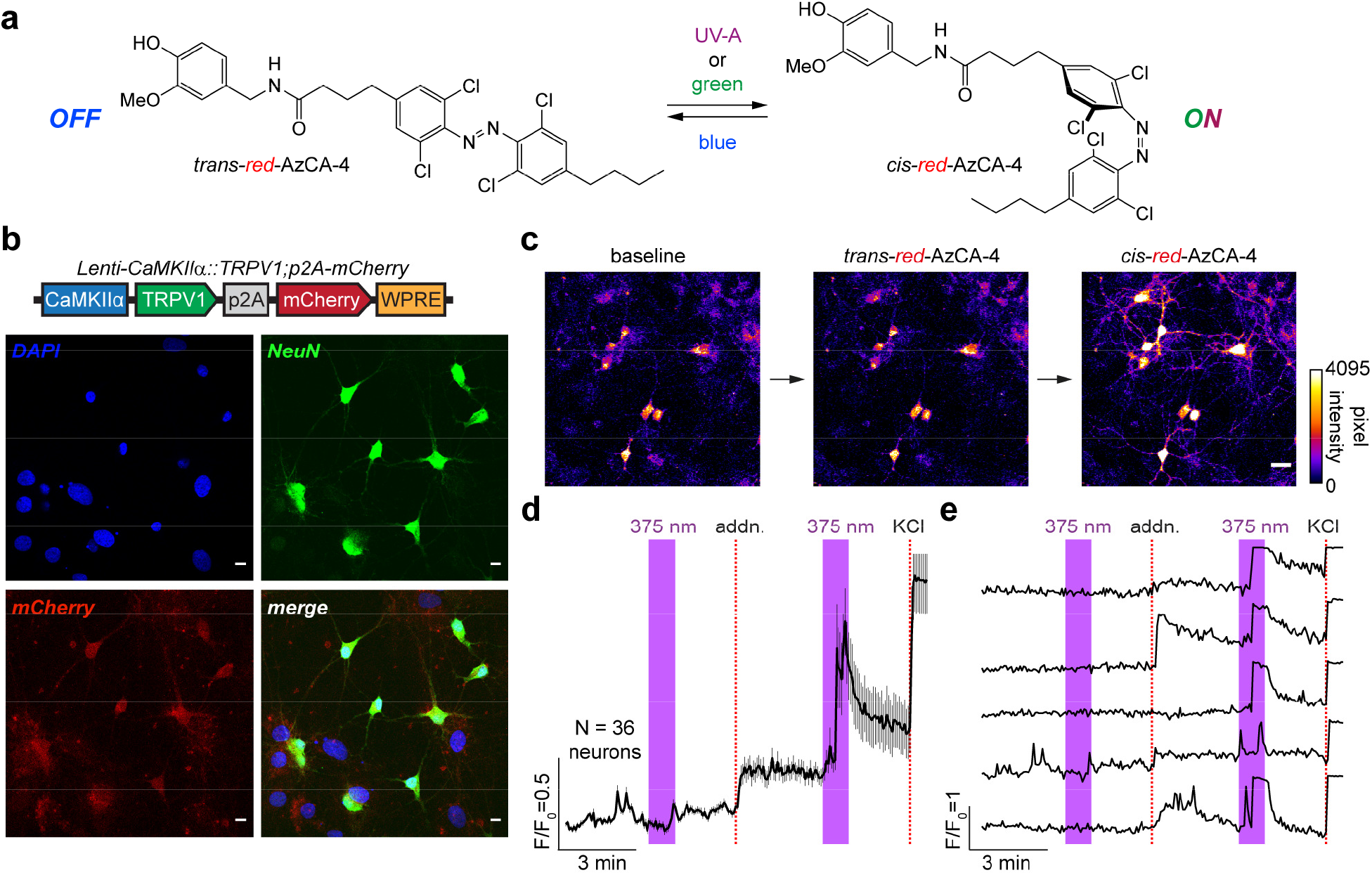
*red*-AzCA-4 enables optical control of TRPV1-infected primary neurons. (**a**) Chemical structure of *red*-AzCA-4, which can be isomerized between its *cis*- and *trans*-isomers with UV-A/green and blue light, respectively. (**b**) Cultured rat hippocampal neurons were infected with a virus encoding for TRPV1 and mCherry under CaMKIIα promoter. Immunofluorescence imaging showed that neurons (NeuN, green) expressed mCherry (red). (**c-e**) Fluorescent Ca^2+^ imaging of Fluo-4-loaded neurons showed that *red*-AzCA-4 (20 nM) stimulated [Ca^2+^]_i_. Displayed as (**c**) representative images, (**d**) average normalized Ca^2+^ levels over multiple cells (N = 36 neurons, 2 experiments, Error bars = mean ± S.E.M.); and (**e**) representative Ca^2+^ traces from five individual neurons.

To sensitize neurons to CAP and *red*-AzCA-4, we utilized a lentivirus carrying the TRPV1 transgene under the excitatory neural promoter calmodulin kinase II α-subunit (CaMKIIα). TRPV1 expression was linked to a red fluorescent protein mCherry by a post-transcriptional cleavage linker p2A^24^ (Lenti-CaMKIIα::TRPV1-p2A-mCherry)^25^ (**Fig. 1b**). In primary rat neurons, lentiviral transduction affected nearly 100% of the cells as marked by colocalization of mCherry expression with neuronal marker NeuN.^26^ To test the activity of *red*-AzCA-4 on transduced neurons, the culture was incubated with the Ca^2+^-sensitive dye Fluo-4-AM, and the intracellular Ca^2+^ concentration ([Ca^2+^]_i_) was monitored using confocal fluorescence imaging (**Fig. 1c**). For these experiments, *red*-AzCA-4 photoswitching was triggered using the 375 nm (UV-A) quenching laser to enable simultaneous imaging of the green Fluo-4-AM fluorescence. No fluorescence increase was observed on addition of the *trans*-compound, however irradiation with a 375 nm laser caused a rapid increase in Fluo-4 fluorescence, demonstrating that *cis*-*red*-AzCA-4 increased the [Ca^2+^]_i_ (**Fig. 1d-e**). In control experiments, addition of CAP also caused an increase in [Ca^2+^]_i_ (**Supplementary Fig. 1a,b**). Additionally, UV-A irradiation alone, or a vehicle addition (0.1% DMSO in buffer) +/− irradiation did not affect [Ca^2+^]_i_ (**Supplementary Fig. 1c,d**). These results indicate that the *cis*-isomer of *red*-AzCA-4 activates TRPV1-more strongly than *trans*. This is consistent with the results obtained in the previous studies,^18,21^ and sets the stage for the *in vivo* application of *red*-AzCA-4.

### Developing a multifunctional fiber for *in vivo* photopharmacology

Fibers capable of optical stimulation alongside chemical/virus delivery were produced *via* thermal drawing of macroscale preforms. The fibers contained a polymer optical waveguide and two microfluidic channels, which were fabricated out of polycarbonate (PC) and cyclic-olefin-copolymer (COC) (**Fig. 2c**, top). These materials possess similar T_g_ (PC, T_g_ = 150 °C; COC, T_g_ = 158°C), making them suitable for co-drawing. By tuning the preform feeding and fiber drawing speeds, the cross section of the preform (3.7×2.6 cm^2^) was reduced to 480×340 μm^2^ in the final fiber (**Fig. 2c**, middle, **Supplementary Fig. 2**). Connectorization of the microfluidic channels to ethyl vinyl acetate (EVA) tubing and the PC/COC waveguide to a ceramic optical ferrule (**Supplementary Fig. 3**) afforded the final implant with light and fluid-delivery capabilities (**Fig. 2c**, bottom). The final device was miniature (2.5×2×0.5 cm^3^) and lightweight (0.82±0.06 g), which permitted unconstrained movement of mice following implantation.

**Figure 2.**
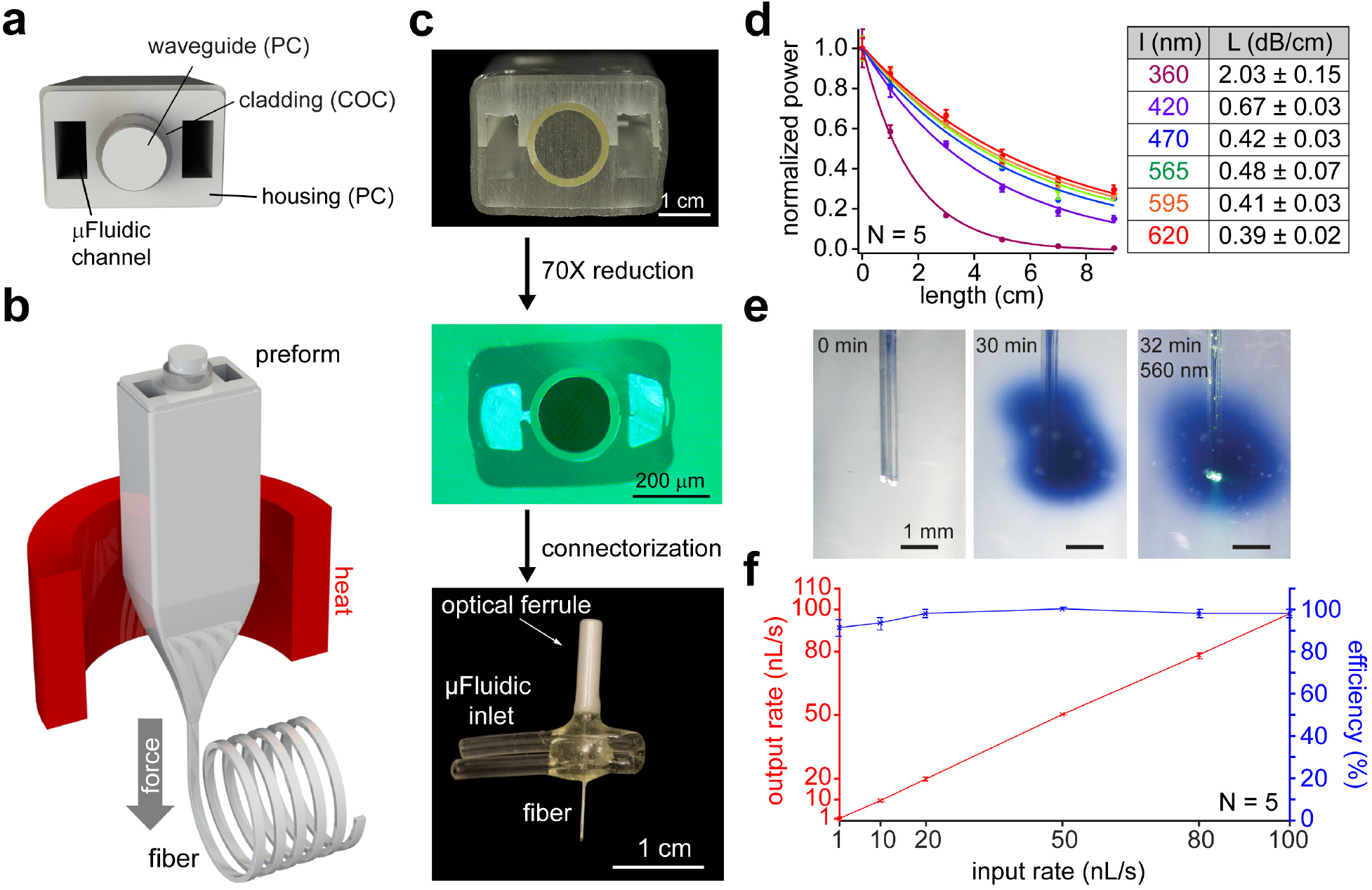
Design and fabrication of a multifunctional fiber-based implant by thermal drawing. (**a**) Cross-section schematic of implants with two μfluidic channels and optical waveguide. (**b**) Schematic representing the thermal drawing process. (**c**) Cross-section of the preforms for before thermal drawing (top). Cross-section of fibers after thermal drawing (middle). Fully connectorized device (bottom). (**d**) Optical loss calculation of the PC/COC optical waveguide across the visible spectrum (N = 5). (**e**) Injection of tryptan-blue dye (2 μL over 30 min) into a phantom brain (0.6% agarose gel). 565 nm LED irradiation emerges from the device tip. (**f**) Output injection rate and injection efficiency of the microfluidic channel of 1 cm long multifunctional fiber-based neural implant (N = 5). Error bars = mean ± S.E.M.

We characterized the optical waveguide using fiber-coupled LEDs across the UV-visible range by measuring the optical loss over different lengths of fiber (**Fig. 2d**, left). Optical losses across the visible range spanned 0.67±0.03, 0.42±0.03, 0.48±0.07, 0.41±0.03, 0.39±0.02 dB/cm for 420, 470, 565, 595, and 620 nm wavelengths, respectively (**Fig. 2d**, right). These values are comparable to those previously obtained in thermally drawn polymer fibers [1.2-2.7 dB/cm].^27–30^ This makes our device suitable for controlling photoswitchable ligands with light across the visible spectrum.^22^ Due to the absorbance of PC used as the waveguide core in the UV-A range, the optical loss at 365 nm was 2.03±0.15 dB/cm. We then tested the microfluidic channels by injecting tryptan-blue dye into a “phantom brain”, consisting of 0.6% agarose gel. No leaks were observed along the fiber lengths, and the fluid delivery did not interfere with optical illumination (**Fig. 2e**). Furthermore, fluid return rates were >80% at injection speeds ranging between 1-100 nL/s (**Fig. 2f**). Taken together, these results demonstrate that our fibers can concurrently deliver both optical and chemical (fluid) stimuli.

### *In vivo* photopharmacology of midbrain dopaminergic circuits

To test the utility of our fibers for delivering and optically actuating photoswitchable ligands to deep brain regions *in vivo*, we implanted our devices into the VTA in mice. The VTA is known as a dopaminergic hub involved in reward signaling and reinforcement, and its dopaminergic projections to the nucleus accumbens (NAc) and medial prefrontal cortex (mPFC) are well characterized (**Fig. 3a**).^31–33^ The fibers were used to deliver Lenti-CaMKIIα::TRPV1-p2A-mCherry virus (1.5 μL, >10^10^ particles/mL) into the VTA during the implantation surgery (**Fig. 3b**). This one-step surgery reduces the tissue damage that may occur during multiple surgeries, and also ensures colocalization of the expression profile with the optical and pharmacological interrogation volume.^28^ Six to eight weeks post-implantation, we confirmed TRPV1 expression and the fiber position in the VTA (**Fig. 3c, Supplementary Fig. 4**).

**Figure 3.**
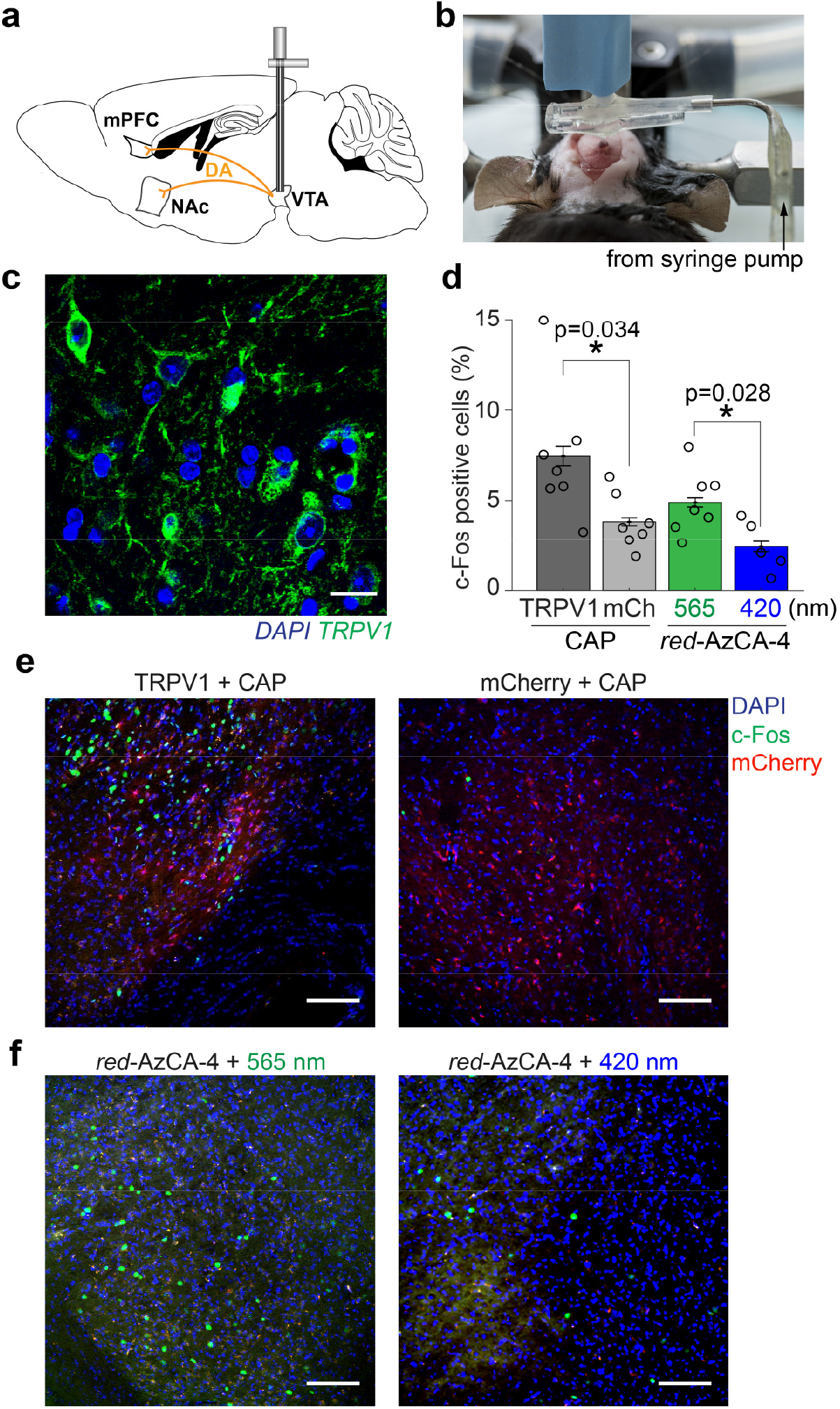
*in vivo* chemogenetic stimulation of midbrain dopaminergic circuits. (**a**) Schematic of VTA dopamine projections to the mPFC and NAc. (**b**) Image during craniotomy during simultaneous implantation and virus injection. (**c**) 6 weeks post-injection, Immunofluorescence imaging using an anti-TRPV1 antibody confirmed TRPV1 expression (green) in the VTA. Co-stained with the DAPI nuclear marker (blue). Scale bar = 10 μm. (**d**) Quantification of c-Fos expression in the VTA following CAP or over multiple animals (TRPV1 + CAP, N=7; mCherry, N=7; *red*-AzCA-4 + 565nm light, N=7; *red*-AzCA-4 + 565nm light, N=5. (**e**) Injection of CAP (10 μM) caused an increase in c-Fos expression (green nuclei) in the TRPV1+ VTA neurons (top left panel) when compared to (top right panel). (**f**) Injection of *red*-AzCA-4. Scale bar = 100 μm. Error bars = mean ± S.E.M.

Next, we tested the ability of CAP and *red*-AzCA-4 to activate transduced TRPV1-expressing VTA neurons *in vivo* by quantifying the expression of c-Fos.^34^ This immediate early gene serves as a marker of neuronal activity, as its expression and translocation to the nucleus is upregulated in recently-activated neurons.^35^ A separate cohort of mice received a Lenti-CamKIIα::mCherry virus as a negative control. CAP or *red*-AzCa-4 were injected through the microfluidic channels of the implanted fibers over 5 min concurrently with illumination. We showed that CAP injection (10 μM) into TRPV1-expressing mice (N = 7) caused an increase in c-Fos expression in the VTA when compared to those only expressing mCherry (N = 7) (**Fig. 3d,e**). The locations of the c-Fos expressing cells correlated with the region displaying mCherry fluorescence. Next, *red*-AzCA-4 (1 μM) was injected under either 420 nm or 565 nm illumination. We observed a greater c-Fos expression in the VTA in the presence of green light (N = 7) compared to blue light (N = 5), indicating that *cis-red*-AzCA-4 activates VTA neurons more strongly than the *trans*-isomer (**Fig. 3d,f**). c-Fos immunofluorescence imaging in the NAc and mPFC also revealed greater c-Fos expression in response to *cis*-*red*-AzCA-4 compared to *trans-red*-AzCA-4 (**Supplementary Fig. 5a,b**). We did not observe an increase in c-Fos expression in TRPV1-expressing mice injected with a vehicle control (0.1% DMSO), demonstrating that the process of infusion itself did not affect c-Fos expression (**Supplementary Fig. 6**). Consistent with prior work, these results demonstrate that multifunctional fibers are capable of delivering both viruses, chemicals, and light to deep brain regions. Notably, both CAP and *red*-AzCA-4 activate sensitized neurons, where *red*-AzCA-4 affords an additional level of optical control. As such, our approach is suitable for photopharmacological manipulation of neural circuits *in vivo*.

### Optical control of place preference using *red*-AzCa-4

Activation of VTA dopaminergic neurons is associated with rewarding/salient stimuli, and is known to drive behavioral preference in mice.^36^ To test the ability of *red*-AzCA-4 to activate the mesolimbic pathway in a light-dependent manner, we performed a three-chamber conditioned place preference (CPP) assay.^37^ Mice were implanted with multifunctional fibers and concomitantly injected with a Lenti-CaMKIIα::TRPV1-p2A-mCherry into the VTA, and the CPP test was performed six to eight weeks post-implantation. Mice injected with Lenti-CaMKIIα::mCherry served as negative controls. The first day of the CPP protocol began with a 45 min pre-test during which each mouse could freely explore a three-chamber arena. The left and right chambers were distinguishable by patterns on the floor and walls (holes vs. bars) (**Fig. 4a**). On the second and third conditioning days, the mice were confined to a single chamber where they received unique chemical and optical stimulation. Between mice, the stimuli were randomized with respect to the chamber. The fourth day consisted of a post-test where each mouse could explore the entire box. The relative time spent on each chamber was compared between day 1 (pre-test) and day 4 (post-test) to determine the preference of the mice for the two stimuli. A cohort of mice was conditioned to compare CAP *vs.* vehicle control, while another received *red*-AzCA-4 injections with either blue or green light (**Fig. 4b**). In the pre-test, TRPV1-expressing mice displayed no preference for either chamber (N = 18) (**Fig. 4c,d**). However, in the post-test, mice spent more time in the chamber in which they received a CAP injection (10 μM) compared to the vehicle control (0.2% DMSO) (N = 7) (**Fig.4c,d**). Similarly, mice injected with *red*-AzCa-4 (1 μM) preferred the chamber in which they received green (*cis*-*red*-AzCA-4) compared to blue illumination (*trans*-*red*-AzCA-4) (**Fig. 4c,d**). Mice expressing mCherry alone did not develop a preference for CAP over the vehicle control (N = 8) (**Fig. 4e**), indicating that exogenous TRPV1 expression in the VTA is necessary to drive the preference. These results are consistent with our *in vitro* studies and c-Fos quantification experiments, where *cis*-*red*-AzCA-4 activates TRPV1 more potently. These findings confirm that *red*-AzCA-4 combined with the multifunctional fibers enables optical control behavior in freely moving mice.

**Figure 4.**
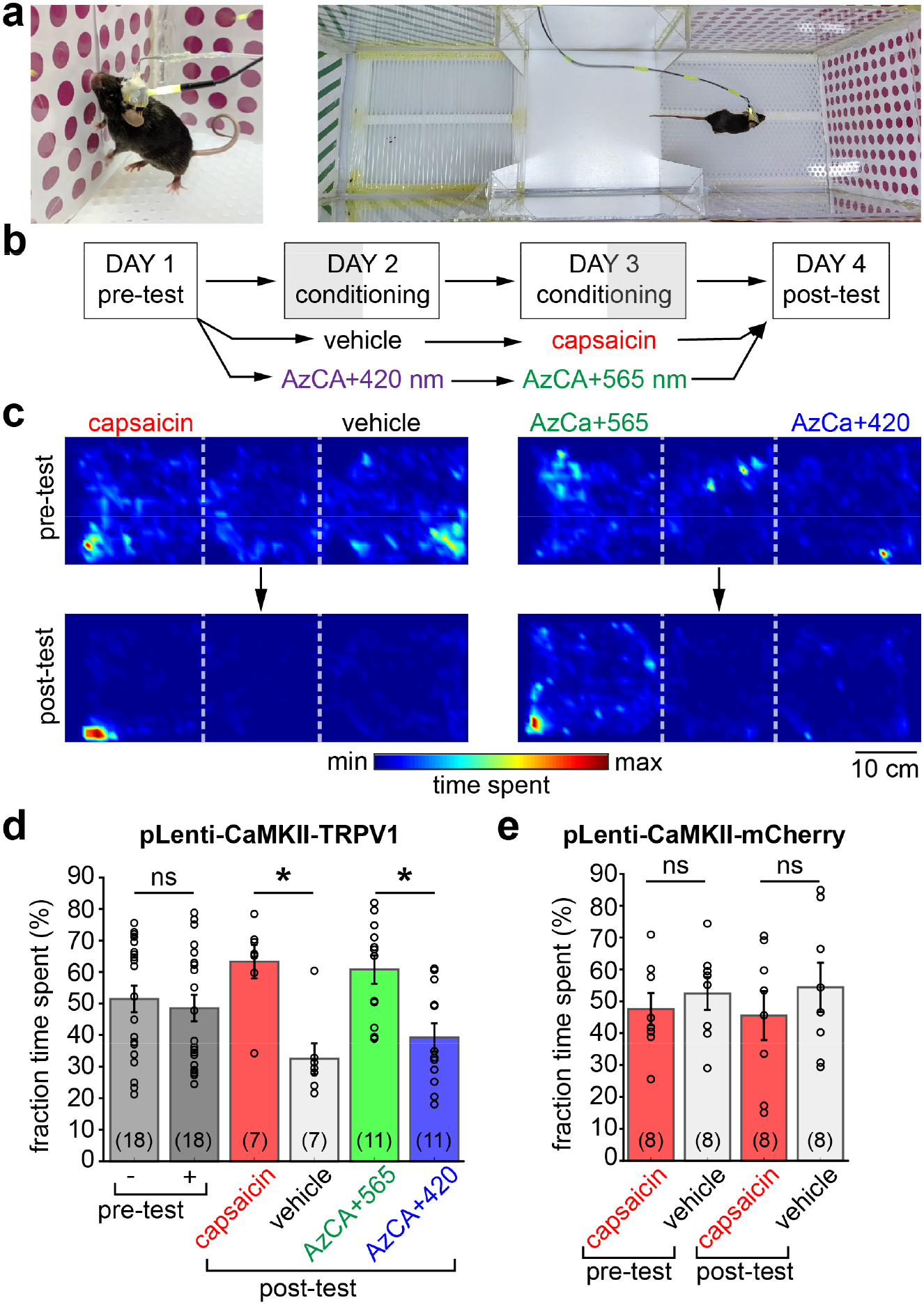
*red*-AzCA-4 enables light-dependent control of reward behavior in freely moving animals. (**a**) Awake and freely moving mice were connected to a microfluidic pump and LEDs and subject to a CPP test. (**b**) Timeline of the 4 day CPP test. (**c**) Heatmaps depicting position in the box for a representative mouse during the pretest (top) and after conditioning (post-test, bottom) with CAP/vehicle (left) or *red*-AzCA-4 + 420/565 nm irradiation (right). (**d**) TRPV1-expressing mice did not have a preference for either chamber during the pre-test day (N = 18). Mice preferred CAP over a vehicle control on the post-test day after conditioning (N = 7). Mice preferred the chamber where they received *cis*-*red*-AzCA-4 (green light) over *trans*-*red*-AzCA-4 (blue light) after their conditioning (N = 11). (**e**) Mice which only expressed mCherry and not TRPV1 did not develop a preference for the CAP stimulated chamber. Error bars = mean ± S.E.M. *P < 0.05, ns = not significant.

## DISCUSSION

Since the first azobenzene-based photoswitches were employed to reversibly activate ion channels,^38–40^ photopharmacology has evolved to the point where an optical switch can be placed on virtually any component of the cellular signaling machinery. Although chemical photoswitches have been extensively applied to *in vitro* and in vertebrates,^9^ their use in the neuroscience community has lagged behind primarily due to challenges in applying these probes to mammalian models. Bringing the chemical flexibility of photopharmacology to the neuroscience community is therefore an important step in understanding the molecular mechanisms underlying behavior.

In this study, we demonstrated that photopharmacological manipulation of ion channels affords optical control of neural circuits sufficient to modulate behavior. Our *in vivo* studies show that green light potentiates the effect of photoswitchable compound *red*-AzCA-4 by generating the more potent *cis*-isomer.^18^ Compared to recent applications of *in vivo* photopharmacology in mice which required a separate cannula and waveguide for probe and light delivery,^15–17^ our multi-functional device can deliver transgenes, ligands, and light to deep brain regions such as the VTA. We anticipate this will increase the reproducibility of behavioral experiments reliant on photopharmacological manipulation of ion channels. Furthermore, thermal drawing permits integration of additional functional features to the implants, including electrophysiological recording. These may facilitate direct correlation of photopharmacological ion channel manipulation to the dynamics of local circuits.

This proof-of-principle study relied on genetic manipulation to introduce the mammalian receptor TRPV1 in the VTA to achieve photopharmacological control of neurons. Our approach is, however, immediately translatable to photoswitchable ligands targeting endogenous proteins. This will enable optical manipulation of neural circuits without the need for genetic editing. For example, *red*-AzCA-4 could target the naturally expressed TRPV1 in the peripheral nervous system, while other photoswitchable ligands could affect cannabinoid receptors in the brain.^41^ Transgene-free optical manipulation of neural circuits is desirable for model organisms beyond mice, where genetic tools are not as readily available. This would also be of significant importance for the development of potential therapeutic applications, where genetic manipulation still remains a significant barrier.

The advent of optogenetics revolutionized how neuroscientists dissect neural circuitry, yet it remains limited to manipulation of electroactive cells via triggering ion transport through microbial proteins. Photopharmacology empowered by integrated multifunctional fibers will permit direct manipulation of endogenous receptors within living organisms. This will facilitate the studies of the roles of specific neurotransmitters and effector pathways in driving the neural circuits that dictate behavior, physiology, and pathology.

## METHODS

### Cell culture media and solutions

#### Dissection solution

Contains (in mM): 160 NaCl, 5 KCl, 0.5 MgSO_4_, 3 CaCl_2_, 5 HEPES, 5.5 glucose with 2.0 mg/L phenol red. The pH was adjusted to 7.4 with NaOH, sterile filtered and aliquoted.

#### Enzymatic solution

Contains dissection solution (10 mL), L-cysteine (2 mg), EDTA (25 mM, 200 μL aliquot), CaCl_2_ (100 mM, 100 μL aliquot), NaOH (1 Nm, 30 μL), DNAse (0.05 mg/mL, Sigma, DN25), papain (100 μL, Sigma, P3125).

#### Serum media

Contains minimum essential medium (MEM, 500 mL, with Earle’s salts, without glutamine), Glutamax (5 mL, Gibco), fetal bovine serum (FBS, 25 mL), MITO+ serum extender (1 mL, Corning, 35506, diluted 3x in H_2_O). Sterile filtered and aliquoted.

#### Inactivation solution

Contains serum media (10 mL), bovine albumin (25 mg), soybean trypsin inhibitor (25 mg, Gibco, 17075-029), DNase I (0.05 mg/mL, Sigma, DN25).

#### Neurobasal A+ media

Contains Neurobasal-A medium (500 mL, Gibco), B-27 supplement (20 mL, Gibco), Glutamax (5 mL, Gibco), FBS (15 mL). Sterile filtered and aliquoted.

#### FUDR inhibitor

Contains uridine (5 mg/mL), 5-fluorodeoxyuridine (2 mg/mL) in MEM (with Earle’s salts, without glutamine). Sterile filtered and aliquoted.

#### Imaging buffer

Contains (in mM): 115 NaCl, 1.2 CaCl_2_, 1.2 MgCl_2_, 1.2 K_2_HPO_4_, 20 HEPES, 20 D-glucose, adjusted to pH 7.4 with NaOH. Sterile filtered and aliquoted.

### Primary rat hippocampal culture

Hippocampal culture experiments were performed in accordance with the IACUC protocol IP00002121 (Optical tools for controlling and recording neurotransmission). The day before the culture, glass-bottom 8-chamber μ-slides (Ibidi, 80827) were coated with poly-L-lysine (PLL, 50 μL per well) and incubated at 37 °C for 25 min. The chambers were washed with phosphate-buffered saline (PBS, 1x, pH 7.4) and dried for 25 min at 37 °C. The chambers were then coated with mouse laminin (50 μL per well, ≈15 ng/mL in PBS, Corning, 354232) and incubated overnight at 37 °C. Neonatal wild-type Sprague Dawley rats (Charles River, Crl:CD, 8-12 pups, P0-P1) were anesthetized on ice and quickly euthanized by decapitation. The hippocampi were quickly dissected in cold dissection solution and then transferred to sterile filtered enzymatic solution (10 mL) in which they were incubated at 37 °C for 25 min. The hippocampi were then transferred to sterile filtered inactivation solution (10 mL) at room temperature for 2 min. All but ≈1 mL of the media was removed by aspiration and the neurons were gently triturated with a 200 μL pipette until a homogeneous cloudy solution was achieved. The dissociated cells were diluted in serum media (1 mL per hippocampus) and then counted (30 μL cells w/ 10 μL tryptan blue). The cells were plated into the PLL/laminin-coated lab-tecs (200,000 cells/chamber) and cultured at 37 °C in 5% CO_2_. 36-48 h later, the media was exchanged for Neurobasal A+ media (220 μL/chamber) and 30 μL FUDR inhibitor solution was added to each well. Neurons were infected with lentivirus (1:2250× dilution of >10^10^ particles/mL concentrated virus in Neurobasal A+ media) at DIV5. Every 3 days the media was exchanged for fresh Neurobasal A+ media (250 μL/chamber).

### Fiber fabrication

The neural implant was fabricated by using the thermal drawing process on a macroscopic template (preform). To fabricate the preform, we rolled several cyclic olefin copolymer (COC 6015; TOPAS) and polycarbonate sheets (PC; Ajedium films) onto a 12.7 mm PC rod (McMaster) to reach a 16.4 mm diameter assembly, and consolidated under vacuum at 175 °C for ≈30 min, forming a single solid COC/PC preform. Two PC slabs (35.45×15×200 mm; McMaster) were machined to have one semi-circular channel (16.5 mm diameter; 200 mm length) and two rectangular grooves (6.35X6X200 mm) were machine on each side of the semi-circular channel. Two aluminum (Al) slabs (McMaster) were milled to form 11.75×5.75×200 mm^3^ bars, around which several Fluorinated ethylene propylene (FEP) films (McMaster) were rolled to form a 12×6×200 mm^3^ spacer, which was placed inside the rectangular groove and sandwiched between the PC slabs prior to consolidation.

The COC/PC preform was then consolidated in a hot press at 175 °C for 1 h at 1000 psi. Finally, additional PC sheets were wrapped around the overall assembly to provide additional support, and the whole assembly was again consolidated under vacuum at 175 °C for 30 min. The resulting preform was then drawn at 300 °C using a custom-built fiber drawing tower as described in previous work.^27,28^

### Fiber connectorization

The Connectorization process is depicted in **Supplementary Fig. 3**. Drawn fiber was cut to length (4 cm for implantation devices) and one end was coated with epoxy (Devcon, 14250) to protect the polycarbonate (PC) waveguide. The outer PC housing was etched away from the top 1.5 cm of the fiber by submerging the tip in CH_2_Cl_2_ for 2 min. After removal of the epoxy tip, the cyclic-olefin copolymer (COC)/PC waveguide was glued (Devcon epoxy, 14250) into a ceramic ferrule (ThorLabs, CF270) and left to dry overnight. The microfluidic channels were each opened below the ferrule using a scalpel, and the fiber was inserted through two pieces of ethyl vinyl acetate (EVA) tubing (0.03” ID, 0.09” OD, McMaster-Carr, 1883T2), with the core of each tube overlapping the openings for the microfluidic channels. One end of each tube and the region in contact with the fiber was sealed with epoxy (Devcon, 14250) and the ferrule was polished using fiber polishing film (ThorLabs) as described by the manufacturer.

### Microinjection and optical stimulation

Fibers were connected to a Nanofil syringe (10 μL, World Precision Instruments) using a home-built connector consisting of a Nanofil beveled needle (33G, World Precision Instruments, NF33BV), EVA tubing (0.03” ID, 0.09” OD, McMaster-Carr, 1883T2) and dispensing needle (19G, McMaster-Carr, straight: 75165A554, or 90°: 75165A66). The individual components were assembled and sealed with 5-min epoxy (Devcon, 14250). Injections were controlled using a syringe pump (World Precision Instruments, Micro4, UMC4) and controller (World Precision Instruments, UMP3).

Optical stimulation was generated using fiber-coupled LEDs (Thorlabs, M420F2, M565F3) driven by a LED driver (Thorlabs, LEDD1B, 0.6 A current limit) and a waveform generator (Agilent 33500B). LEDs were coupled to the ferrule on the device using a rotary joint optical fiber (Thorlabs, RJPSF2) and a ceramic mating sleeve (Thorlabs, ADAF1).

### Microfluidic characterization - phantom brain injection and injection efficiency

Agarose gel (Sigma) was dissolved in distilled H_2_O (0.6% wt/vol) with gentle microwave heating, poured into a 2 mL glass vial and let cool overnight. The fiber tip was inserted into the agarose gel and the injection was driven using a syringe pump (WPI Micro4, UMC4) and controller (WPI, UMP3), via a Hamilton syringe and EVA tubing (McMaster, #1883T1). Tryptan blue (3 μL) was injected at a speed of 100 nL/min. Images were captured using a Nikon D610 digital single-lens reflex camera and a 100 mm macro lens (Tokina). RAW images were processed in Adobe Photoshop, where corrections were applied only to increase sharpness and color contrast.

To further characterize the microfluidic capabilities of the neural implant, the microfluidic inlet of the device was connected to the injection apparatus described above (Nanofil, syringe pump and controller). We injected PBS (9 μL) at a target injection rate of 1, 10, 20, 50, 80 and 100 nL/s through the microfluidic channel, and injection output was measured by weight. Output injection rate was calculated by dividing the injection output by the time required to inject it. Injection efficiency was calculated by comparing input and output injection rate.

### Optical loss calculation

To calculate the optical loss, fibers (11 cm in length) were etched and connected to a ferrule (ThorLabs, CF270) as described above. The fiber was connected to fiber-coupled LEDs (Thorlabs, M365FP1, M420F2, M470F3, M565F3, M595F2, M625F2) using a rotary joint optical fiber (Thorlabs, RJPSF2) and a metal interconnect (Thorlabs, ADAF2). The absolute output power was measured using digital power meter (Thorlabs, PM100D) with a photodiode power sensor (Thorlabs, S121C). 2 cm sections of fiber were then removed and the power at each length was measured down to 1 cm. The recorded power values at each length were normalized to the power at the shortest length. This procedure was repeated for multiple fibers, and the normalized powers were averaged and plotted (**Fig. 2d**). The decibel loss was calculated across each cut section as:

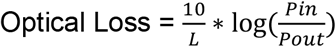

in dB/cm, where L = fiber length in cm; P_in_ = input power in mW; P_out_ = output power in mW. The values across the 11 cm segments of each fiber were averaged, and the error was calculated as ±S.E.M.

### Lentivirus packaging

Plasmids for lentivirus packaging were amplified by transformation into stellar competent cells (ClonTech) and extracted with endotoxin-free maxi-prep DNA extraction kit (Qiagen). Lenti-X HEK 293T cells (ClonTech) were used for producing lentiviral particles, and were maintained in 10 cm petridish (Corning) in DMEM with 10% FBS (Thermofisher) without antibiotic for >3 passages before transfection. Cells were transfected with the 3rd generation lentivirus and helper plasmids when the cells reached 80≈90% confluence. Before transfection, HEK cells were incubated in 6 mL fresh and prewarmed DMEM with 10% FBS for 1 h. Plasmids and linear polyethylenimine (PEI; MW = 25000, Polysciences) were premixed and incubated at room temperature for 25 min in the following amounts per plate: 2 mL opti-MEM (GIBCO), 3 μg pRTR2; 3 μg pVSVG; 9 μg p4.1R; 15 μg pLenti-CaMKII-TRPV1-p2A-mCherry-WPRE or pLenti-CaMKII-mCherry-WPRE; 90 mg PEI and incubated in room temperature for 25 min. The transfection mixture were added to each plate (2 mL) and incubated at 37 °C for 6-18 h. The culture medium was then exchanged with 10 mL fresh and prewarmed DMEM with 10% FBS. 48-72 h later, the culture medium was collected in 50 mL centrifuge tubes. The cell debris were removed by centrifugation at 3000 rpm for 5 min at 4 °C. The lentiviral medium was filtered with a 0.45 μm filter and stored in 4 °C before concentration. The approximate titer of lentivirus was evaluated using a lentiviral titration kit (Lenti-X GoStix Plus; ClonTech). The lentiviral medium was transferred to 50 mL centrifuge tubes, and 7 mL 10% sucrose in PBS (Corning) was slowly added into the bottom of each tube. Lentivirus was pelleted by centrifugation at 11,000xg for 4 h at 4 °C.^42^ After removal of the supernatant and drying of the centrifuge tubes, the lentivirus pellets were resuspended in PBS at 0.1% the original volume of the lentiviral medium. The titer of lentivirus was approximated by a lentiviral titration kit (Lenti-X GoStix Plus; ClonTech) with serial dilutions. The concentrated lentivirus was aliquoted and stored at −80 °C freezer until use.

### Stereotactic surgery and device implantation

All stereotactic surgeries were performed in accordance with the IACUC protocol 0118-003-21 (Implantation of Neural Recording and Stimulation Devices into Adult Mice and Rats). Mice were housed in a normal 12 h light/dark cycle and fed a standard rodent chow diet. In brief, 8 week old male BL6C/57 mice (Jackson, BL6C/57, #000664) were anesthetized with 1-2% isoflurane and placed on a heat pad in a stereotaxic head frame (Kopf) and immediately injected subcutaneously with buprenorphine-slow release (ZooPharm, 1.0 mg/kg). After creation of a midline incision along the scalp, a craniotomy was performed using a rotary tool (Dremel Micro 8050) and a carbon steel burr (Heisinger, 19007-05). Implants were flushed with sterile PBS, fastened to the stereotaxic cannula holder and connected to a Hamilton syringe (10 μL, World Precision Instruments). Concentrated lentivirus (>10^9^ transducing units/mL for LentiCaMKIIα::TRPV1-p2A-mCherry or Lenti-CaMKIIα::mCherry) was loaded into a microfluidic channel and the device was lowered into the skull (coordinates from bregma, in mm: ML: ±0.5, DV: −4.35, AP: −3.3). The virus solution (2 μL total) was slowly injected (flow rate: 75 nL/min) while the device was cemented to the skull using first C&B-Metabond adhesive cement (Parkell) and then dental cement (Jet Set-4) to encase the device. After completion of the injection, the device remained attached to the Hamilton syringe for at least 10 min before being detached. The incision was then closed with sutures, and the mouse was given a subcutaneous injection of carprofen (5 mg/kg) and sterile Ringer’s solution (0.6 mL) prior to recovery on a heat pad. For 3 days post-implantaion, mice were singly housed and closely monitored for signs of overall health. They were given Carprofen injections (0.6 mL, 0.25 mg/mL in sterile Ringer’s solution) as necessary. After recovery, mice remained singly housed and were not used for experiments if they experienced symptoms of decreased health.

### Injection for c-Fos quantification

Six to eight weeks post device implantation (above), mice were anesthetized with 2% isoflurane and the microfluidic channels were flushed with compound sterile PBS 3x with a Hamilton syringe and 26G needle. The device was connected via mini-EVA tubing (McMaster, 1883T1) to a Hamilton syringe that had been pre-filled with compound solution. Compound solution (3 μL total volume) was injected (600 nL/min), and for *red*-AzCA-4 injections, 420 or 565 nm irradiation was applied (2 Hz, 50% duty cycle) for 40 min (as described above). The animal was kept under anesthesia for a total of 90 min after the beginning of the injection to allow for c-Fos expression, and then deeply anesthetized by Fatal-plus injection (0.075 mL of 97.5 mg/mL solution in NaCl) followed by a standard transcardial perfusion using PBS (50 mL) followed by PFA (Electron Microscopy Sciences, 50 mL, 4% in PBS).

### Immunofluorescence staining

For immunofluorescence in cultured neurons, the coverslips were fixed with 4% PFA (Electron Microscopy Sciences, 190107) in PBS buffer for 20 min at room temperature and then washed out for 5 min with PBS. Coverslips were permeabilized with 0.2% Triton X-100 (Fisher Scientific, BP151-100) for 1 min and then washed with PBS for (3×5 min washes). The coverslips were then transferred to a new 24-well plate (Corning) and incubated in blocking buffer (0.02% Triton X100 + 5% normal donkey serum [NDS] in PBS) for 1 h at room temperature on an orbital shaker (100 rpm). The coverslips were then incubated in primary antibody solution (1:300 dilution anti-NeuN antibody [Abcam ab177487], 0.02% Triton X100, 5% NDS in PBS). The plate was sealed with parafilm (Pechiney plastic) and incubated overnight (12-14 h) in the dark at room temperature on the orbital shaker. The coverslips were washed with PBS (3×5 min washes) on the shaker at room temperature, and then transferred to the secondary antibody solution (1:1000 donkey anti-rabbit 488 secondary [Life Technologies A21206], 0.02% Triton X100, 5% NDS) and incubated for 1 h in the dark at room temperature on the shaker. The coverslips were again washed with PBS (3×5 min washes) and then incubated in the dark with a 2-[4-(aminoiminomethyl)phenyl]-1H-Indole-6-carboximidamide hydrochloride (DAPI) solution (1:50000 dilution, from 5 mg/mL DMSO stock in PBS) for 10 min on the shaker. The coverslips were washed with Sudan Black (Sigma Aldrich, MKCG8561) for 3×5 min washes (0.2 mg/mL in 70% EtOH) and washed with deionized water (2×5 min washes). Coverslips were then mounted on a microscope slide (VWR® Superfrost® Plus Micro Slide, 75×25×1 mm, 48311-703). The coverslips were covered with mounting solution (Fluoromount™), and stored in the dark at room temperature overnight before imaging.

For fixed brain slice immunofluorescence, mice were euthanized by standard transcardial perfusion using 4% PFA (PFA, Electron Microscopy Sciences) in PBS. After extraction, the brains were soaked in 4% PFA in the dark overnight at 4 °C. The brains were washed with PBS (3×15 mL for 15 min), sectioned appropriately and coronal or sagittal slices (60 μm thickness) were prepared with vibratome (Leica VT1000S) with FEATHER® razor blade (Electron Microscopy Sciences, 72002) in ice-cold PBS buffer. The slices were stored in the dark in PBS buffer at 4 °C until staining. The slices were transferred to a Netwell™ insert (Corning, 3478) and then incubated in permeabilization/blocking buffer (0.3% Triton X100 + 3% normal donkey serum [NDS] in PBS) in a standard 12-well plate (Cellstar, 665180) for 1 h at room temperature in the dark on an orbital shaker (100 rpm). The slices were then transferred to the primary antibody solution (2 mL total volume, 1:1000 dilution primary antibody, 3% NDS in PBS). The plate was sealed with parafilm (Pechiney plastic) and incubated overnight (12-14 h) in the dark at room temperature on the shaker. The slices were washed with PBS (3×2 mL, 20 min each) on the shaker in the dark at room temperature, and then transferred to the secondary antibody solution (2 mL total volume in PBS) and incubated for 2 h in the dark at room temperature on the shaker. The slices were again washed with PBS (3×2 mL, 20 min each) and then incubated in the dark with a DAPI solution (1:50000 dilution, from 5 mg/mL DMSO stock in PBS, 2 mL total volume) for 45 min on the shaker. The slices were again washed with PBS (2×2 mL, 20 min each) and then mounted on a cover slide (VWR^®^ Superfrost^®^ Plus Micro Slide, 75×25×1 mm, 48311-703). The slices were covered with mounting solution (Fluoromount™), covered with a coverglass (VWR^®^, 24×60 mm, 16004-096) and then stored in the dark at room temperature overnight.

### Confocal microscopy

Live-cell fluorescent Ca^2+^ imaging in cultured hippocampal neurons (DIV 10-14) was performed on a dual scanner Olympus Fluoview 1200 with a 20× objective (Olympus) at 37 °C in 5% CO_2_. Cells were first loaded with Fluo-4-AM (2 μM, ThermoFisher, F14201) in Neurobasal A+ media for 30 min, and then washed twice with imaging buffer (250 μL). Fluo-4 was excited at 488 nm at low laser power (<3%), and the emission collected from 500–550 nm. Photoswitching was generated by 375 nm laser irradiation at 100% laser power, and triggered using the quench function in the Olympus software. The fluorescence intensity data was extracted in ImageJ (NIH) and processed/plotted in MATLAB.

Fluorescence images in fixed and mounted sagittal brain slices were acquired on an Olympus Fluoview FV1000 laser-scanning confocal microscope with a 20× objective (oil, NA=0.85), and 60× (oil, NA=1.42) objective. For images of each brain region, serial z-stack images with a step size of 4 μm were acquired. The center of the frame was positioned at: VTA - ML: ±0.48 mm, DV: −4.4 mm, AP: −3.3 mm; mPFC - ML: ±0.36 mm, DV: −2.5 mm, AP: +1.5 mm; Nac - ML: ±0.72 mm, DV: −3.9 mm, AP: −1.25 mm.^43^ Coordinates were determined for each region using the Alan Brain Atlas. DAPI was excited using a 405 nm laser with emission light collected between 425-460 nm. Alexa Fluor 488 imaging was excited using a 473 nm laser with emission light collected between 485-515 nm. mCherry was visualized using a 559 nm laser with collected emission light between 575-675 nm.

Following imaging, the data was analyzed with a custom MATLAB script (available on request) and ImageJ (NIH), which directly processed output .oib files without user intervention to generate the z-stacks, and quantify the percentage of c-Fos positive cells. Statistical significance was assessed in Matlab using a two-sample t-test. All imaging and analysis were blinded with respect to the experimental conditions.

### Conditioned place preference

Conditioned place preference (CPP) experiments were performed in a home-built three chamber box with dimensions 21×25, 21×20, and 21×25 cm for the left, center, and right chambers, respectively. The base was fabricated out of white Polyethylene sheets (McMaster, 8752K315) to facilitate cleaning between trials. The left chamber had a white plastic floor with punched holes (2.38 mm diameter, 25% open area) (McMaster #9293T67) and walls with magenta polka-dot pattern. The center chamber had a flat Teflon floor (McMaster) with no wall pattern. The right chamber had a floor made of plastic bars (1/8 inch diameter, 7.62 mm spacing) (McMaster, 8658K47) and diagonal green bars on the walls. The walls and doors to separate the chambers were fabricated out of transparent polycarbonate (McMaster). A webcam (Logitech), microinjector pump (World Precision Instruments, Micro4, UMC4), and the rotary joint of the optical waveguide (Thorlabs, RJPSF2) were mounted directly above the center chamber. Experiments were performed in a room with normal fluorescent lighting and the light intensity was kept constant across the box and consistent throughout all experimental days. Video recording of all trial days (days 1-4) were recorded using the webcam software (Logitech). The absolute position of the mouse in the box was determined using a custom python script (available on request), and the percent time spent in each chamber of the box was calculated using MATLAB (script available on request). The first 30 min of the video were used for quantification, and the time in either the holes vs. bars chamber were scored as a percentage. Pre-conditioning was performed for 3 days prior to the behavioral experiments to habituate them to being restrained and attached to optical/fluid delivery cables. Animals were transported to the behavioral space and the microfluidic channels were flushed with 3×10 μL sterile PBS. The experimental paradigm for the 4-day conditioned place preference (CPP) test is as follows:

#### DAY 1, pre-test

The microfluidic channel was flushed with 3×10 μL washes of PBS buffer and the device was connected to an optical fiber and the injection tubing. The mouse was then placed in the center chamber with closed doors. The doors were then removed and the mouse was left to explore the entire box for 45 min.

#### DAY 2, condition 1

Microfluidic channels were flushed with 3×10 μL washes of compound solution (0.1% DMSO or 1 μM *red*-AzCA-4) and the device was connected to the optical fiber and injection tubing. The mouse was placed in either the left or right chamber (the mice were evenly distributed between holes or bars), which was sealed by the transparent door. The trial was initiated on beginning of the injection (3 μL at 600 nL/min) and the mouse was kept in the conditioning chamber for 45 min. For trials with *red*-AzCA-4, 420 nm light was applied (7.6 mW/mm^2^ on average) at 2 Hz at 50% duty cycle. After the trial, the channels were again washed with 3×10 μL sterile PBS before the mice were returned to the animal facility overnight.

#### DAY 3, condition 2

The procedure for day 3 was the same as day 2, except the mouse was placed in opposite chamber. On this day, the mice which previously received the vehicle control received a 10 μM CAP injection. The mice which previously received the *red*-AzCA-4 with blue light received the same dose of *red*-AzCA-4, except with 565 nm irradiation (2 Hz at 50% duty cycle, 8.1 mW/mm^2^ on average).

#### DAY 4, post-test

Identical to the procedure described above for day 1.

### Statistical methods and error reporting

Unless otherwise described, all data is presented as mean ±S.E.M. For Ca^2+^ imaging experiments, n is the number of measurements made (individual cells) and the number of individual experiments is described in the figure caption. Statistical significance was assessed using Matlab (Mathworks). For the comparison between two groups in immunohistochemistry analyses and behavior assays, Student’s two sample t-test were used, with significance threshold placed at *P <0.05.

## ACKNOWLEDGEMENTS

Authors thank Prof. Dirk Trauner for his input on this study. This work was funded in part by National Institute of Neurological Disorders and Stroke (5R01NS086804), the National Institutes of Health (NIH) BRAIN Initiative (1R01MH111872), and the National Science Foundation (NSF) Center for Neurotechnology (EEC-1028725), and the McGovern Institute for Brain Research. This work made use of the MIT MRSEC Shared Experimental Facilities under award number DMR-14-19807 from the NSF. JAF acknowledges the Vollum Institute Fellowship for financial support. MJA is a recipient of the Friends of McGovern Graduate Fellowship.

## AUTHOR CONTRIBUTIONS

JAF and PA conceived and coordinated the study. DBK synthesized the photoswitchable compounds. JAF and GR carried out imaging experiments in cultured neurons. JAF, MJA, and IG fabricated and characterized the multifunctional fibers. YF aided in fiber design. PC and FK produced lentiviral vectors. JAF, MJA, PC, AC, and IG performed immunohistochemical experiments. JAF and MJA performed the behavioral experiments. PC, AC, MJA, and JAF wrote scripts for data analysis. JAF, MJA, and PA wrote the manuscript with input from all co-authors.

## COMPETING INTERESTS

The authors declare no competing interests.

