## Supplementary Information for "*In vivo* photopharmacology enabled by multifunctional fibers"

**SUPPLEMENTARY FIGURES**

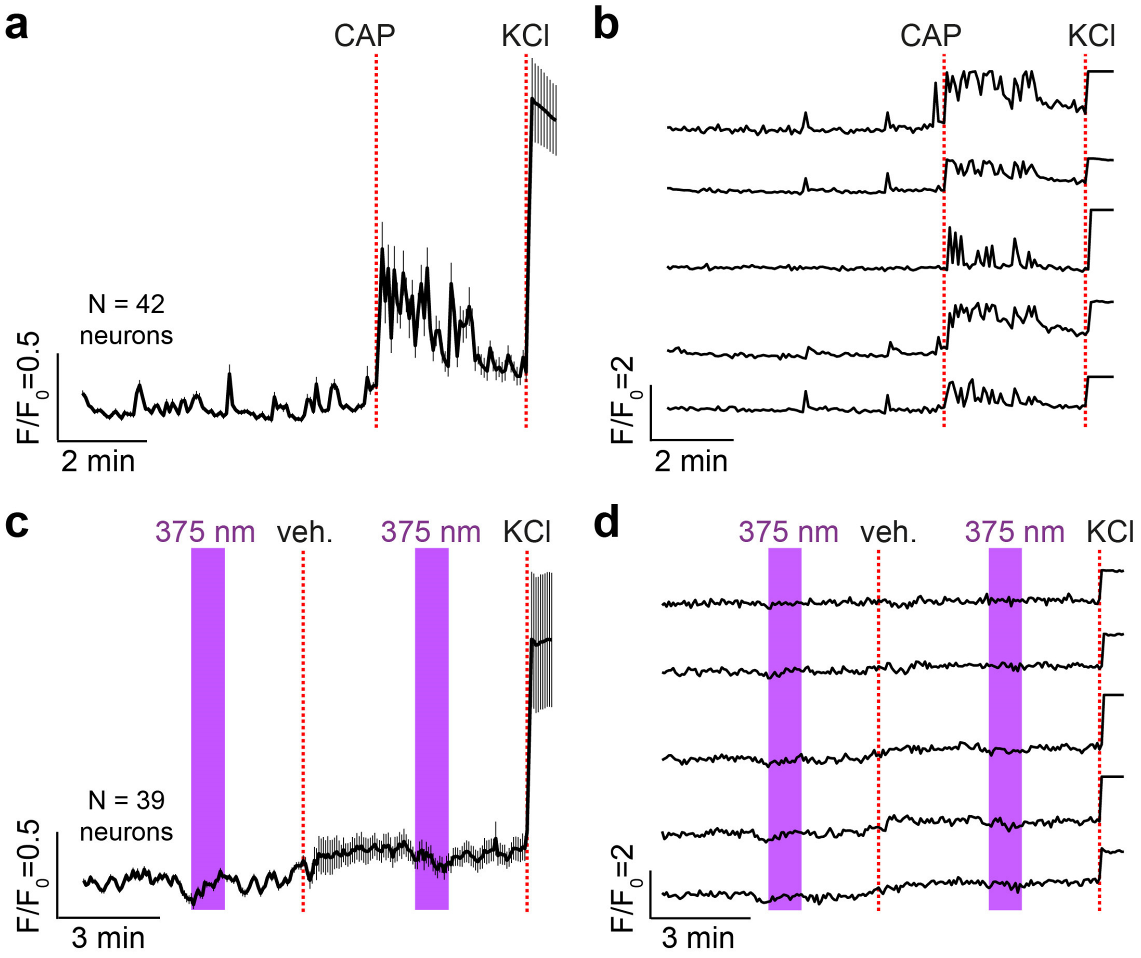


**Supplementary Figure 1 | *red*-AzCA vehicle control experiments in cultured neurons**. Fluo-4 fluorescence was recorded in cultured rat hippocampal neurons that had been transduced with LentiCaMKIIα::TRPV1-p2A-mCherry virus. (**a**,**b**) CAP addition (20 nM) increased intracellular $Ca^{2+}$ levels. Displayed as (**a**) the average normalized fluorescence level (N = 42 neurons from 2 experiments) and (**b**) 5 traces from representative neurons (**c**,**d**) Addition of a vehicle control (0.1% DMSO, addn.) did not affect $Ca^{2+}$ levels before or after 375 nm irradiation. KCl (25 mM) addition still increased $Ca^{2+}$ levels. Displayed as (**c**) the average normalized fluorescence level (N = 39 neurons from 2 experiments) and (**d**) 5 traces from representative neurons. Error bars = mean ± S.E.M.

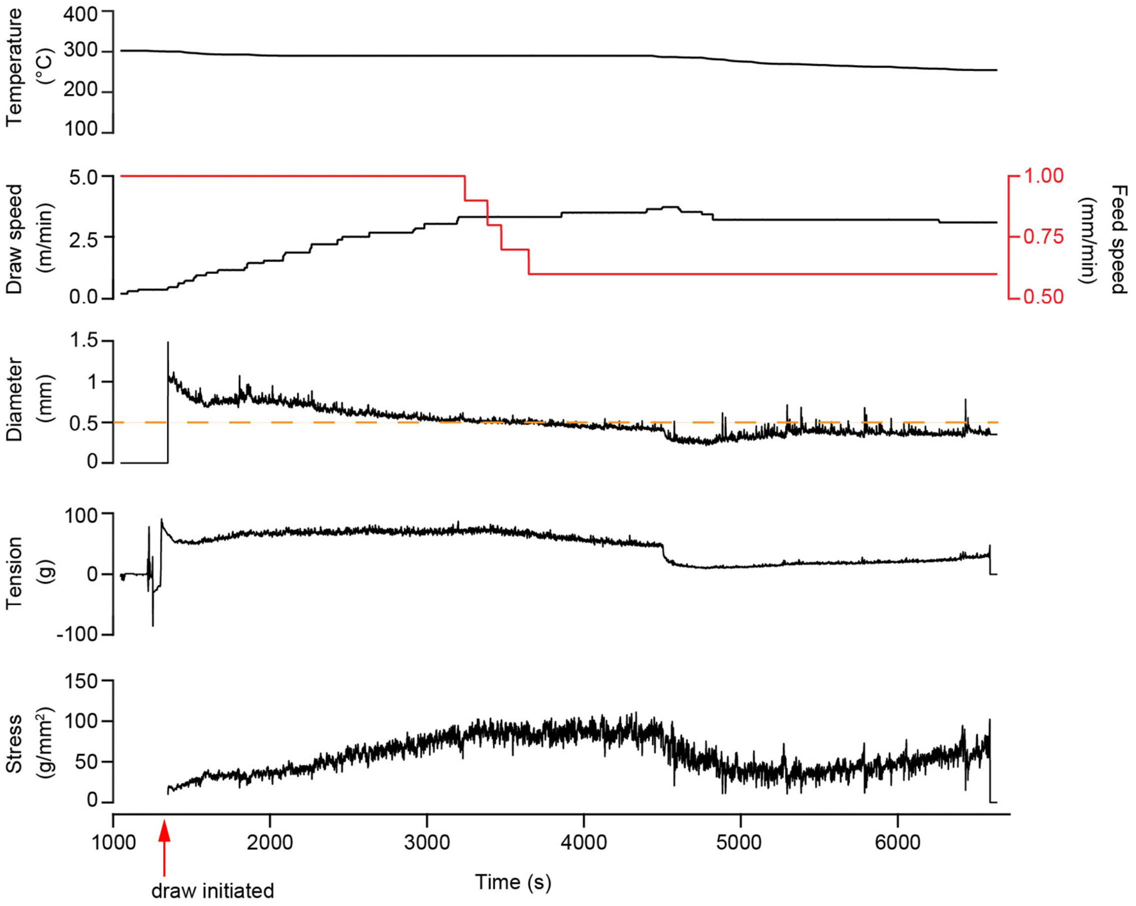


**Supplementary Figure 2 | Drawing parameters and the thermal drawing process.** The thermal drawing process was utilized to draw the macroscopic preform into a microscopic fiber. Displayed are the oven temperature, the draw/feed speeds, the final fiber diameter, and the resulting tension and stress values. The fiber used in this study was taken between t = 5000-6000 s.

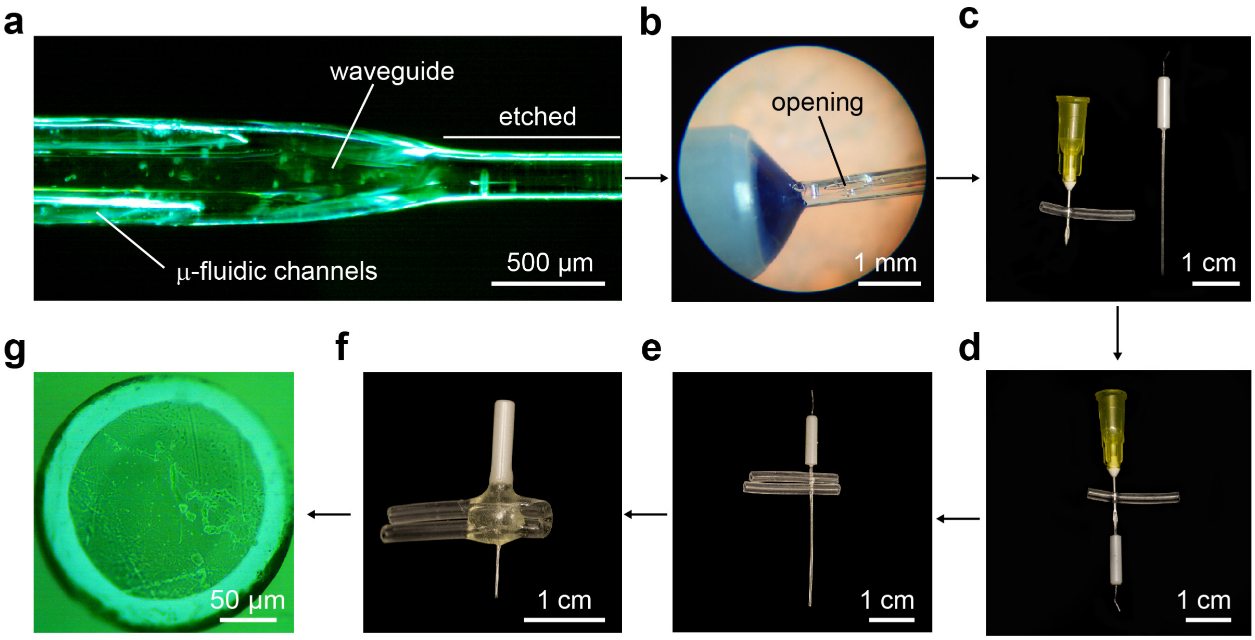


**Supplementary Figure 3 | Fiber Connectorization process.** (**a**) One end of the fiber tip was etched in dichloromethane for insertion into the optical ferrule. (**b**) After gluing into an optical ferrule, the microfluidic channels were manually opened just below the ferrule. (**c**,**d**) A piece of tubing was pierced with a needle, through which the fiber was inserted to slide into the tubing. (**e**) Two pieces of tubing were positioned above the hole for each microfluidic channel. (**f**) The tubing and fiber were sealed with epoxy and (**g**) the ferrule was polished to afford a fully connectorized device (as shown in Fig. 2c).

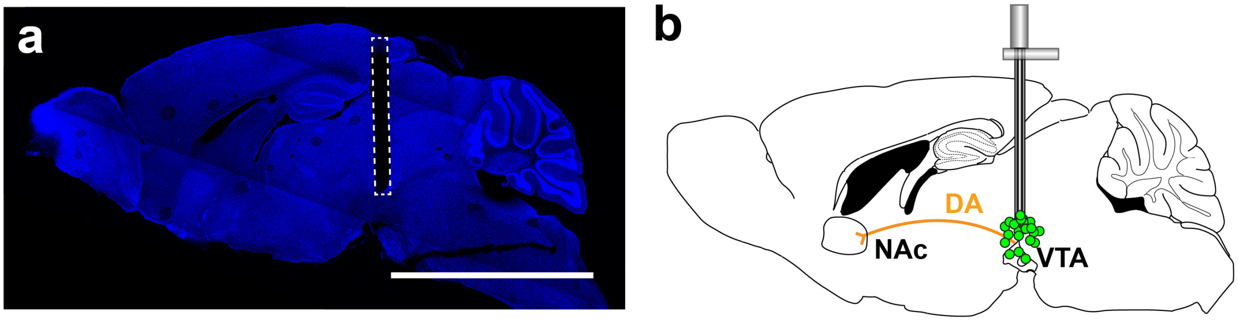


**Supplementary Figure 4 | Implantation locations.** (**a**) A representative mosaic fluorescence micrograph of a mouse sagittal brain slice (ML = 0.5 mm, 60 μm thick) stained with DAPI. The fiber implantation location in the VTA is depicted by the dotted line. (**b**) Schematic of all injection positions for the c-Fos immunofluorescence experiments, depicted as colored dots.

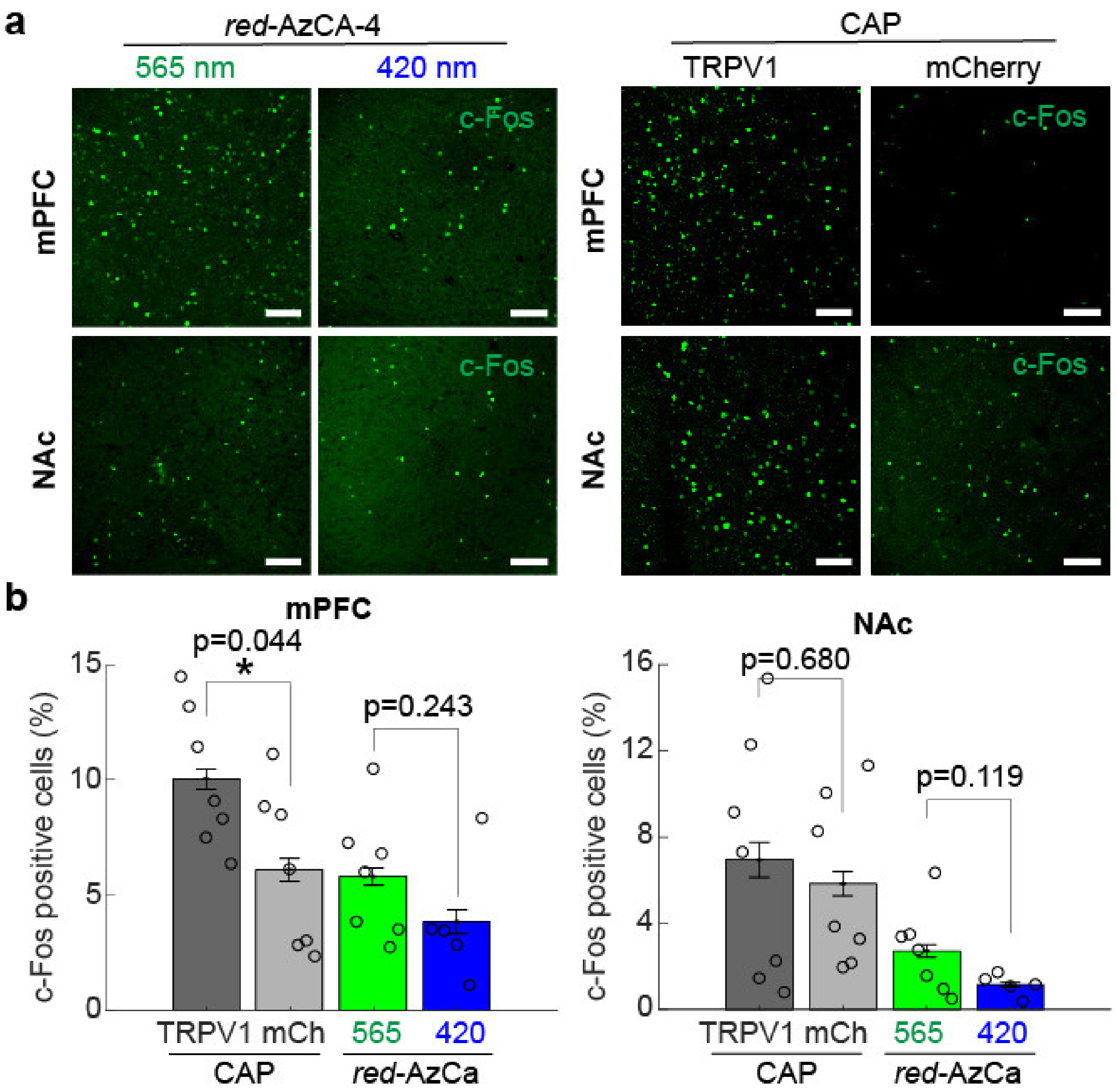


**Supplementary Figure 5 | Chemogenetic control of VTA projections.** *Red-AzCA-4* (3 μL, 1 μM) and CAP (3 μL, 10 μM) injection in anesthetized mice increased c-Fos expression in the NAc and mPFC. For *red*-AzCA-4, c-Fos was further upregulated in the presence of green (565 nm) light (N=7) compared to blue (420 nm) light (N=5). Similarly, CAP injection into mice expressing TRPV1 (N=7) upregulated c-Fos expression compared to control mice expressing mCherry only (N=7). Displayed as (**a**) representative images from fixed VTA slices, and (**b**) a quantification of the % c-Fos positive cells from multiple animals. Error bars = mean ± S.E.M.

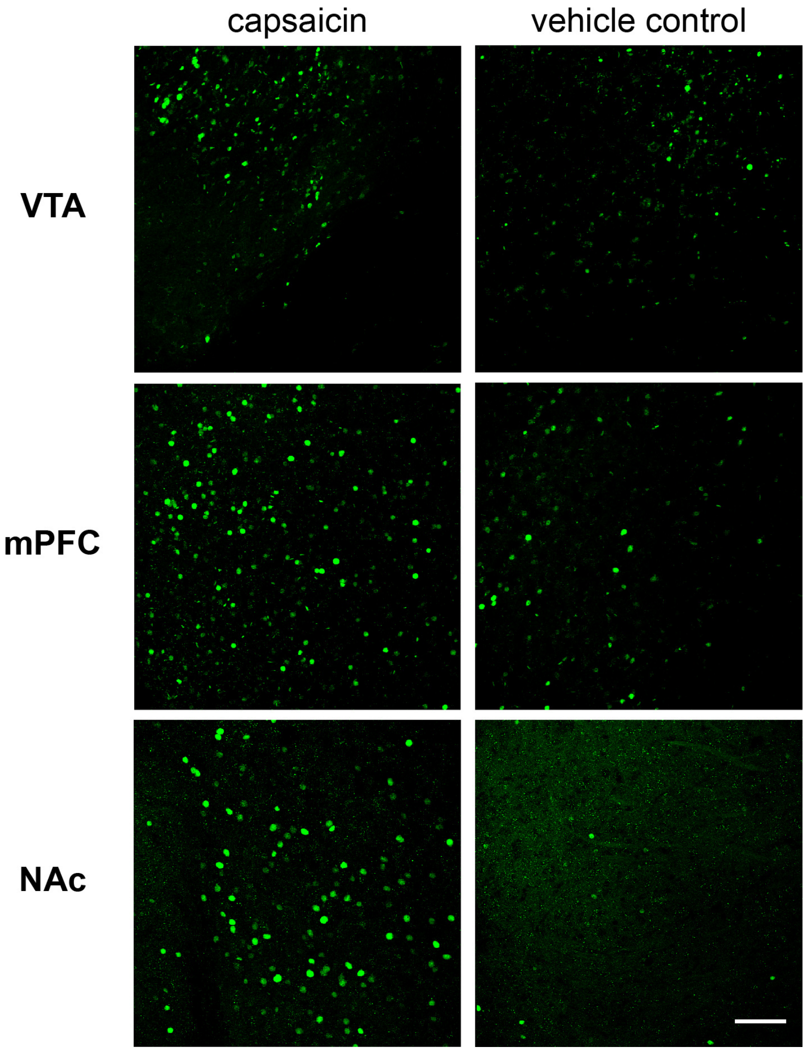


**Supplementary Figure 6 | *in vivo* chemogenetic stimulation vehicle control experiment.** Immunofluorescence imaging in TRPV1-infected mice showed increased nuclear c-Fos expression in the VTA (top), mPFC (middle), and NAc (bottom) after CAP (5 μM, 1.5 μL, left) when compared to a vehicle control (0.1% DMSO, 1.5 μL, right). Scale bar = 100 μm.

**Chemical reagents and synthesis**

All reactions were magnetically stirred under inert gas ($N_2$) atmosphere using standard Schlenk techniques. Glassware was evacuated and dried by heating with a heat-gun (set to 550 °C). Drying over $Na_2SO_4$ implies stirring with excess anhydrous salt for 3-5 min followed by filtration through a glass frit and rinsing of the filter cake with additional solvent. Cannulas and syringes which were used for transferring reagents or solvents were flooded with inert gas (3×) before use. Purification by column chromatography was performed under elevated pressure (flash column chromatography) on Geduran© Si60 silica gel (40-63 μm) from Merck KGaA. After flash column chromatography, the concentrated fractions were filtered once through a glass frit. Silica gel $F_{254}$ TLC plates from Merck KGaA were used for monitoring reactions, analyzing fractions of column chromatography and measuring $R_f$ values. Drying via lyophilization or freeze-drying refers to freezing of the respective sample in liquid nitrogen followed by evacuating the containing flask with high vacuum (<1 mbar) and slow thawing to room temperature. Reaction yields refer to spectroscopically pure isolated amounts of compounds. NMR-spectra were acquired on a Bruker Avance III HD 400 with Cryo-head (400 MHz for $^{1}H$ and 101 MHz for $^{13}C$ spectroscopy). High-resolution mass spectra were recorded on a Thermo Finnigan LTQ FT (ESI: electrospray ionization).

Unless otherwise specified, chemicals were purchased from *Sigma Aldrich*, *Fisher Scientific*, *TCI Europe*, *Chempur*, *Alfa Aesar* or *Acros Organics*. Solvents purchased in technical grade quality were distilled under reduced pressure and used for purification procedures. Triethylamine ($NEt_3$) was dried by distillation from $CaH_2$. Dry EtOAc was purchased from commercial sources (*Acros Organics*, *Fisher Scientific*) under inert gas atmosphere and over molecular sieves. All other reagents with a purity of >95% were purchased from commercial sources and used without further purification. *Red*-FAAzo-4 was synthesized according to our literature procedure.[1] The large-scale synthesis of red-AzCA-4 is as follows:

### Synthesis of *red*-AzCA-4

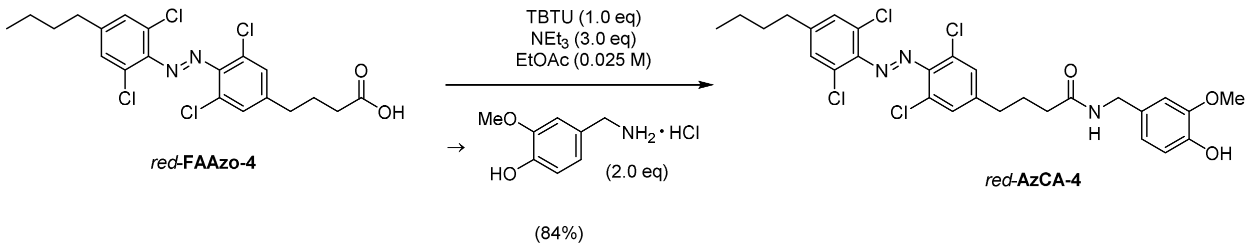


*Red*-FAAzo-4 (700 mg, 1.48 mmol, 1.00 equiv.) was placed in a Schlenk flask and freeze-dried (2 cycles). 2-(1H-Benzotriazole-1-yl)-1,1,3,3-tetramethylaminium tetrafluoroborate (TBTU, 476 mg, 1.48 mmol, 1.00 equiv.) was added and the mixture was dissolved in EtOAc (59.3 mL). After a dropwise addition of $NEt_3$ (0.62 mL), the red solution was stirred for 70 min at room temperature. Vanillylamine hydrochloride (562 mg, 2.96 mmol, 2.00 equiv.) was added in one portion and stirring was continued at room temperature for 2 h. The solution was diluted with EtOAc (250 mL) and washed with 3% $KHSO_4$ (50 mL) and saturated aqueous NaCl (100 mL) followed by drying over $Na_2SO_4$ and removal of the solvent *in vacuo*. Two consecutive column chromatography steps (petane:EtOAc = 1:1) afforded *red*-**AzCA-4** (740 mg, 1.24 mmol, 84%) as a dark red solid. The analytical data is in accordance with those reported in the literature.[1]

**Table 1: Corresponding primary and secondary antibodies for immunohistochemistry**

| Protein Target | Primary Antibody | Secondary Antibody |
|---|---|---|
| NeuN | Anti-NeuN Rabbit mAb; Abcam, #ab177487, lot #: GR3250076-4; 1:300 dilution | Donkey anti-Rabbit IgG (H+L) Highly Cross-Adsorbed Secondary Antibody, Alexa Fluor 488; Invitrogen, #A-21206, lot #: 2045215; 1:1000 dilution |
| c-Fos | Anti-c-Fos (9F6) Rabbit mAb; Cell Signaling Technology, #2250s, lot #: [Ref: 09/2019 Lot:10]; 1:1000 dilution | Donkey anti-Rabbit IgG (H+L) Highly Cross-Adsorbed Secondary Antibody, Alexa Fluor 488; Invitrogen, #A-21206, lot #: 1927937; 1:2000 dilution |
| TRPV1 | Anti-Capsaicin Receptor Antibody, NT; Chemicon®, AB5889, lot #: 3022017; 1:1000 dilution | Donkey anti-Rabbit IgG (H+L) Highly Cross-Adsorbed Secondary Antibody, Alexa Fluor 488; Invitrogen, #A-21206; 1:2000 dilution |
